# Obesity-elicited macrophages shape CD9^hi^ progenitor fate to promote adipose tissue fibrosis and dysfunction

**DOI:** 10.1101/2023.09.26.559540

**Authors:** Haoussa Askia, Sébastien Dussaud, Marina Chaves de Oliveira, Amanda Oliveira, Lucie Poupel, Clémentine Rebière, Melissa Ouhachi, Raoul Manuel, Fatiha Merabtene, Laurent Genser, Judith Aron-Wisnewsky, Martine Moreau, Laurent Yvan-Charvet, Adaliene Ferreira, Thierry Huby, Karine Clément, Geneviève Marcelin, Emmanuel L. Gautier

## Abstract

Obesity is a life-threatening condition characterized by a maladaptive remodeling of the visceral white adipose tissue (vWAT), including fibrosis, that drives vWAT metabolic alterations. We previously identified CD9^hi^ adipose tissue progenitors as the main drivers of vWAT fibrosis in mice and humans. However, how their functions are controlled, especially by macrophages, remains largely unknown. We found that obesity-elicited monocyte-derived macrophages (obeMac) accumulation was considerably elevated in mice prone to obesity-induced vWAT fibrosis. Limiting obeMac build-up decreased the numbers and fibrogenic activation of CD9^hi^ progenitors, leading to decreased vWAT fibrosis and improved glucose homeostasis. In patients with obesity, we identified macrophages similar to mouse obeMacs that were associated with accumulation of CD9^hi^ progenitors in the vWAT and loss of glycemic control. Finally, intercellular communication analysis identified mediators produced by obeMacs that control the fibrogenic potential of CD9^hi^ progenitors. Together, we uncovered an obeMac-CD9^hi^ progenitors axis controlling vWAT fibrosis and dysfunction.

## INTRODUCTION

Obesity sets a state of chronic, metabolic inflammation (or metaflammation), that in turn favors obesity-associated comorbidities onset and progression ^1^. The white adipose tissue (WAT) is a major site where metaflammation arise, leading to the build-up of an inflammatory infiltrate dominated by macrophages ^2^. Macrophages (Macs) locally produce inflammatory mediators, including cytokines, that directly acts on adipocytes to alter their function and impact on whole-body metabolic fitness ^3^. Besides promoting inflammation and altering adipocytes functions, obesity also drives WAT ultrastructural remodeling ^4^. Such maladaptive remodeling, which mainly relies on extracellular matrix (ECM) deposition, also participates to induce and fuel WAT dysfunction ^4^. We previously identified PDGFRα^+^ adipose tissue progenitors with high cell surface levels of the tetraspanin CD9 as critical ECM producers in mice prone to obesity-induced visceral WAT (vWAT) fibrosis and in the vWAT of patients with obesity ^5^. Inflammation is tightly associated with the onset of fibrosis in multiple pathological contexts and *Tlr4* invalidation in hematopoietic cells dampened ECM deposition in the vWAT of obesity-induced fibrosis prone mice ^6^. As *Tlr4* is preferentially expressed by macrophages in the mouse immune system ^7^, this suggest a crosstalk between macrophages and adipose tissue progenitors may operate to induce vWAT fibrosis and metabolic impairments. Yet, such crosstalk has not been uncovered so far nor the nature of the macrophage population involved.

Macrophages are diverse in origins with tissue-resident Macs (trMacs) cohabitating with inflammatory monocyte-derived Macs (moMacs) in pathological conditions ^8^. Diversity in the macrophage pool populating the obese vWAT has been long recognized, with macrophage bearing the CD11c marker accumulating during obesity development ^9^. Here, we set out to further characterize macrophages in the healthy and obese vWAT. We then focused on those infiltrating the vWAT during obesity progression and how they potentially impact vWAT remodeling, especially in the control of adipose tissue fibrosis. We observed that monocyte-derived macrophages accumulation during the course of high-fat diet feeding correlates with the emergence of obesity comorbidities. In a mouse model of obesity-induced adipose tissue fibrosis, we observed that obesity-elicited monocyte-derived macrophages accrual is markedly accelerated. We then showed that these obesity-elicited macrophages favor fibrosis development by increasing the number and the activation of pro-fibrogenic adipose tissue progenitors. In humans, a macrophage subset similar to mouse monocyte-derived macrophages was found to associate with loss of glycemic control and markers of fibrosis in the vWAT. Together, our work underlines a yet unappreciated communication between monocyte-derived macrophages and adipose tissue progenitors in the maladaptive remodeling of the visceral adipose tissue during obesity.

## RESULTS

### Discriminating homeostatic from disease-elicited macrophages in the adipose tissue of obese C57BL6/J mice

We set out to better characterize vWAT adipose tissue macrophages of C57BL6/J mice in order to study their dynamics in the context of diet-induced obesity. Among live CD45^+^ from the stromal vascular fraction (Figure S1A), adipose tissue macrophages are often described as F4/80^+^ CD11b^+^ cells, although those markers are expressed in a variety of myeloid cell types. Our work previously identified CD64 as a more reliable macrophage marker ^7^, which is now used as a standard. Macrophages (F4/80^+^ CD11b^+^ Siglec-F^neg^ CD64^+^), together with eosinophils (F4/80^+^ CD11b^+^ Siglec-F^+^), dendritic cells (F4/80^+^ CD11b^+^ Siglec-F^neg^ CD64^neg^ CD11c^+^ MHC-II^+^) and two undefined populations, fall into the F4/80^+^ CD11b^+^ population in both lean and HFD-fed obese animals (Figure 1A). In obese animals, CD64^+^ Macs appeared heterogenous with only a subset acquiring CD11c expression (Figure 1A) as previously described ^9^. Macs recruited to the obese vWAT most likely fall into this CD11c-expressing fraction, although CD11c did not seem to highlight a clearly definite population. This prompted us to find markers unequivocally discriminating trMacs from those recruited during obesity. Our past experience incited us to look for markers highly expressed on trMacs, rather than recruited macrophages, as trMacs usually express specific markers that makes them more easily recognizable ^7^. Using the Immunological Genome (Immgen) database, we found CD209a, LYVE1, SIGNR1, MRC1, CD301 and CD38 as candidate markers. Among them, CD38, MRC1, CD301 and CD209 were uniformly expressed by trMacs in the vWAT (Figure 1B) and combining those markers distinctly identified trMacs in this depot (Figure 1C). Staining of the CD64^+^ Mac pool of HFD-fed obese mice further pointed to CD38, MRC1 and CD209a as the best candidates to identify trMacs in the vWAT (Figure S1B). Indeed, combinations of those markers revealed two distinct populations, namely CD38^hi^ MRC1^hi^ CD209a^hi^ trMacs and an obesity-elicited CD38^lo^ MRC1^lo^ CD209a^lo^ Mac population (Figure 1D). The later population was also distinctly visible in genetically-obese *Ob*/*Ob* animals (Figure S1C). Noteworthy, this obesity-elicited Mac population couldn’t be well identified using the CD11c marker as CD11c was expressed only a proportion of CD38^lo^ MRC1^lo^ Macs (Figure 1D and Figure S1C).

**Figure 1.**
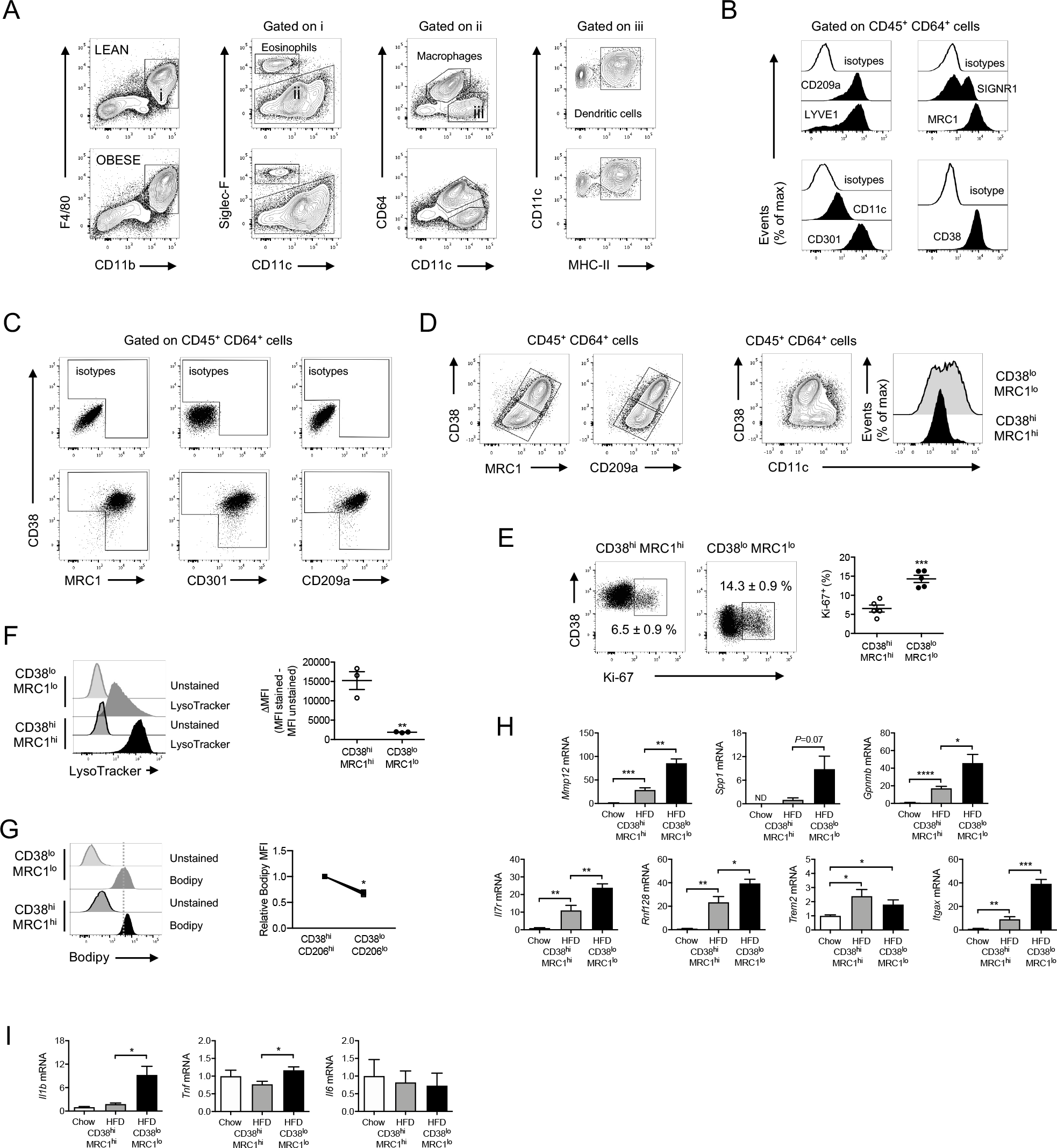
Discriminating homeostatic from disease-elicited macrophages in the adipose tissue of obese C57BL6/J mice. (A) Flow cytometry analysis of myeloid cells in the vWAT of lean and obese mice. (B) Cell surface expression of CD209a, LYVE1, SIGNR1 (CD209b), MRC1, CD11c, CD301 and CD38 on CD64^+^ vWAT macrophages of lean mice. (C) Co-expression of CD38, MRC1, CD301 and CD209a by CD64^+^ vWAT macrophages of lean animals. (D) The expression of CD38, MRC1 and CD209a defines two macrophage subsets in the vWAT of obese mice. (E) Expression of the proliferation marker Ki-67 by CD38^hi^ MRC1^hi^ and CD38^lo^ MRC1^lo^ macrophages in the vWAT of obese mice (n=5 mice). (F) Quantification of lysosomes with Lysotracker staining of CD38^hi^ MRC1^hi^ and CD38^lo^ MRC1^lo^ macrophages in the vWAT of obese mice (n=3 mice). (G) Intracellular lipid levels were determined by Bodipy staining of CD38^hi^ MRC1^hi^ and CD38^lo^ MRC1^lo^ macrophages in the vWAT of obese mice (n=3 mice). (H) RT-qPCR analysis of monocyte-derived macrophage and obesity-associated markers expression in CD38^hi^ MRC1^hi^ and CD38^lo^ MRC1^lo^ macrophages sorted from the vWAT of lean and obese mice (n=3-5 mice per conditions). (I) RT-qPCR analysis of inflammatory cytokines expression in CD38^hi^ MRC1^hi^ and CD38^lo^ MRC1^lo^ macrophages sorted from the vWAT of lean and obese mice (n=3-5 mice per conditions).

Phenotypic characterization of CD38^hi^ MRC1^hi^ Macs revealed greater granularity and size than CD38^lo^ MRC1^lo^ Macs (Figure S1D). Analysis of the Macs proliferative status using Ki67 staining also showed CD38^lo^ MRC1^lo^ cells proliferated 2 to 3 times more than their CD38^hi^ MRC1^hi^ counterparts in HFD-fed mice (Figure 1E) as well as in *Ob*/*Ob* mice (Figure S1E). Then, we found lysosomes (Figure 1F) and lipid droplets (Figure 1G) were more abundant in CD38^hi^ MRC1^hi^ Macs as compared to their CD38^lo^ MRC1^lo^ counterparts, suggesting a stronger lipid catabolic activity. A similar increased lysosomal content was observed in CD38^hi^ MRC1^hi^ Macs of *Ob*/*Ob* mice as compared to CD38^lo^ MRC1^lo^ cells (Figure S1F). Next, we set out to know more about the molecular signature of those Macs subsets, especially the CD38^lo^ MRC1^lo^ population that arise during obesity. Our analysis of a publicly available transcriptomic dataset of the mouse obese vWAT revealed that 30% of the genes induced at the whole tissue level are genes we previously found induced during monocyte to macrophage differentiation *in vivo* ^10^ (Figure S2). Strikingly, more than half of the top 20 induced genes, which include *Mmp12, Gpnmb, Il7r, Rnf128, Trem2* and *Itgax*, were associated with monocyte to macrophage differentiation (Figure S2). This suggests that monocyte-derived macrophage accumulation is a hallmark of the transcriptional changes observed in the obese vWAT, and that this signature might tightly be associated with the presence of obesity-elicited CD38^lo^ MRC1^lo^ macrophages. Analyzing several of the genes cited above revealed they were mostly found in CD38^lo^ MRC1^lo^ Macs, although they were also induced, albeit at lower levels, in CD38^hi^ MRC1^hi^ Macs of obese animals as compared to lean controls (Figure 1H). In addition, CD38^lo^ MRC1^lo^ Macs expressed the pro-inflammatory genes *Tnf* and *Il1b* stronger than CD38^hi^ MRC1^hi^ Macs upon HFD feeding, indicative of a higher activation state (Figure 1I). Overall, using specific adipose trMacs markers, we defined two distinct Mac subsets in the obese vWAT: tissue-resident CD38^hi^ MRC1^hi^ Macs and obesity-elicited CD38^lo^ MRC1^lo^ Macs.

### Macrophage subsets kinetics and metabolic dysfunctions onset over the course of diet-induced obesity in C57BL6/J mice

We then assessed the dynamics of the two Mac subsets defined above during the course of diet-induced obesity. For that purpose, male mice were fed a high-fat diet (HFD) over a 16-week period, and cohorts were analyzed every 4 weeks. Body mass increased overtime due to increased fat mass while lean mass remained stable (Figure 2A). Interestingly, vWAT mass peaked after 8 weeks of HFD, and then lost its ability to further expand over the next 8 weeks of HFD (Figure 2B). Fasting glucose levels slightly increased overtime while insulinemia raised dramatically (data not shown), leading to a progressive increase in the HOMA-IR, an index of insulin resistance (Figure 2C). Notably, we observed the HOMA-IR raised at the time vWAT lost its expandability potential. We next set out to determine whether changes in vWAT Mac subsets content were associated with the onset of metabolic comorbidities. First, we observed that the number of CD38^hi^ MRC1^hi^ Macs in the vWAT increased after 4 weeks of HFD, and then remained stable for the last 12 weeks of diet (Figure 2D). By contrast, obesity-elicited CD38^lo^ MRC1^lo^ cell numbers raised overtime, with a marked increased between 8 and 12 weeks of diet (Figure 2D). We identified the 8-week time point, signing the start of metabolic alterations, as the pivotal time where CD38^lo^ MRC1^lo^ equaled CD38^hi^ MRC1^hi^ Macs in numbers (Figure 2D). After 8 weeks of HFD, metabolic markers strongly worsened in animals while CD38^lo^ MRC1^lo^ Macs started to markedly outnumber CD38^hi^ MRC1^hi^ (Figure 2D). When macrophage density was assessed (*e*.*g*. total cell number divided by tissue weight), we observed CD38^hi^ MRC1^hi^ Macs increased during the first 4 weeks when the vWAT experienced a phase of healthy growth characterized by a very low accumulation of CD38^lo^ MRC1^lo^ cells (Figure 2E). Then, CD38^hi^ MRC1^hi^ cells density did not further increase while obesity-elicited CD38^lo^ MRC1^lo^ Macs started to accumulate to eventually account for the vast majority of ATMs after 12 weeks of diet (Figure 2E). Thus, the two Mac subsets undergo different kinetics over the course of HFD-induced obesity. When vWAT Mac density was plotted together with vWAT mass, it highlighted a clear relationship between CD38^lo^ MRC1^lo^ Macs accumulation and the loss of vWAT expandability capacity (Figure 2F). Similar observations were made in *Ob*/*Ob* mice regarding Mac subsets kinetics and vWAT growth (Figure S3A-B). Interestingly, hepatic parameters (liver mass and signs of steatosis, plasma transaminase activity) also increased in HFD-fed animals at the time vWAT stopped growing (Figure S4A), and vWAT CD38^lo^ MRC1^lo^ Macs, but not CD38^hi^ MRC1^hi^ Macs, strongly correlated with liver mass (Figure S4B). Such correlation also held true in *Ob*/*Ob* animals (Figure S3C). Overall, obesity-elicited CD38^lo^ MRC1^lo^ Macs accumulation in the vWAT is timely correlated to the onset and further aggravation of obesity metabolic comorbidities.

**Figure 2.**
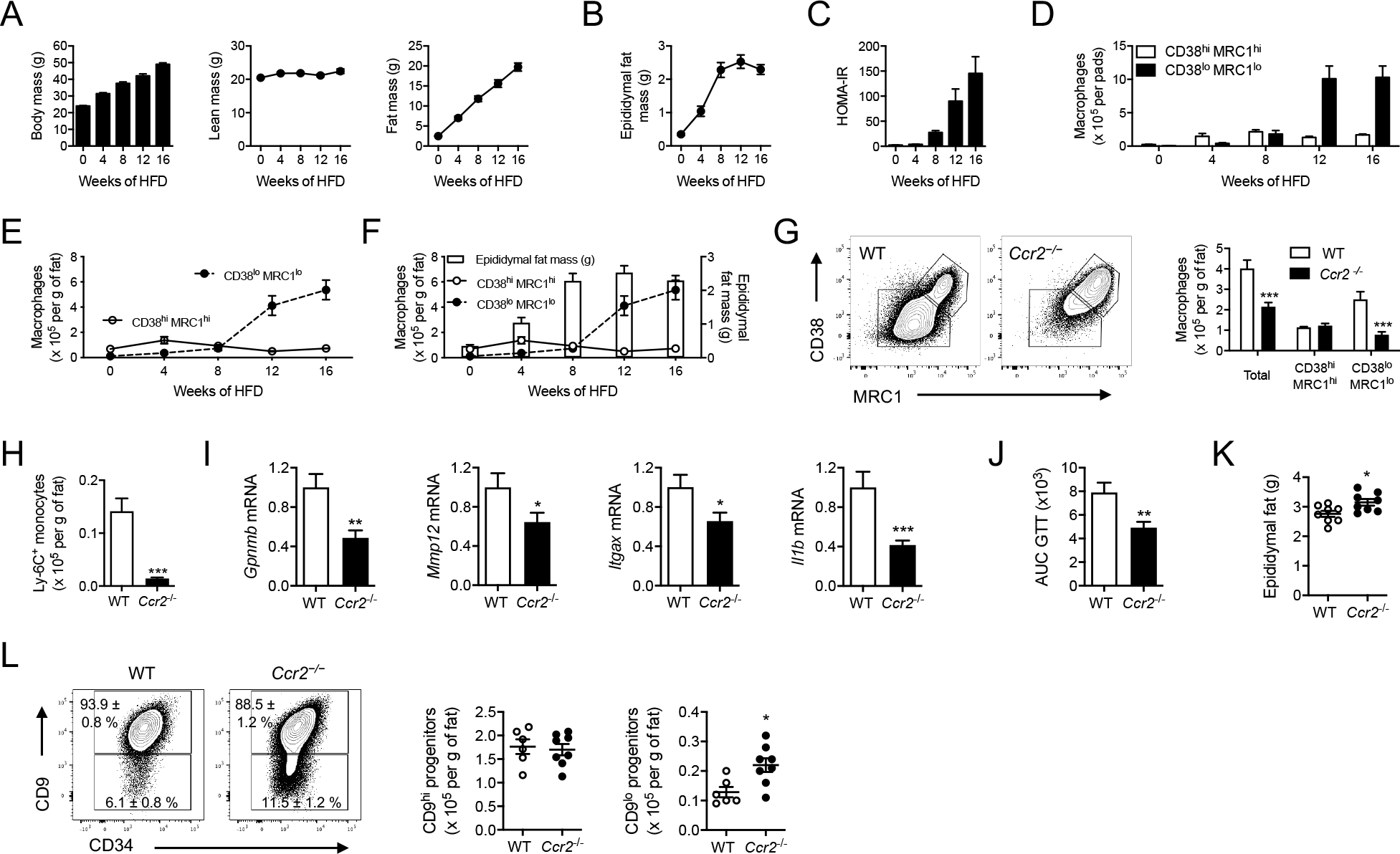
Obesity-elicited CD38^lo^ MRC1^lo^ macrophages accumulation decreases pro-adipogenic progenitors and impairs visceral adipose tissue expansion in HFD-fed C57BL6/J mice. (A) Body mass and body composition (lean and fat mass) before and after 4, 8, 12 and 16 weeks of HFD feeding (n=4-5 mice per time point). (B) Visceral (epididymal) fat mass before and after 4, 8, 12 and 16 weeks of HFD feeding (n=4-5 mice per time point). (C) HOMA-IR index before and after 4, 8, 12 and 16 weeks of HFD feeding (n=4-5 mice per time point). (D) Macrophage subsets numbers in the visceral adipose tissue (pool of the 2 epidydimal fat pads) before and after 4, 8, 12 and 16 weeks of HFD feeding (n=4-5 mice per time point). (E) Macrophage subsets density (number of cells per g of tissue) in the visceral adipose tissue before and after 4, 8, 12 and 16 weeks of HFD feeding (n=4-5 mice per time point). (F) Macrophage subsets density (lines) and visceral (epididymal) fat mass (bar) over the time course of HFD-induced obesity (n=4-5 mice per time point). (G) Flow cytometry analysis and quantification of macrophage subsets density in the visceral adipose tissue of *Ccr2*^−/−^ and wild-type animals after 12 weeks of HFD (n=12-13 mice per group). (H) Quantification of Ly-6C^+^ monocytes density in the visceral adipose tissue of *Ccr2*^−/−^ and wild-type animals after 12 weeks of HFD (n=12-13 mice per group). (I) RT-qPCR analysis of monocyte-derived macrophages markers expression in the visceral adipose tissue of *Ccr2*^−/−^ and wild-type animals after 12 weeks of HFD (n=12-13 mice per group). (J) Area under the curve (AUC) quantification of a glucose tolerance test (GTT) performed on *Ccr2*^−/−^ and wild-type animals after 12 weeks of HFD (n=7 mice per group). (K) Visceral (epididymal) fat mass in *Ccr2*^−/−^ and wild-type animals after 12 weeks of HFD (n=8 mice per group). (L) Flow cytometry analysis and quantification of adipose progenitor subsets density in the visceral adipose tissue of *Ccr2*^−/−^ and wild-type animals after 12 weeks of HFD (n=6-8 mice per group).

### Obesity-elicited CD38^lo^ MRC1^lo^ Macs derive from monocytes and their accumulation limits healthy vWAT expansion in HFD-fed C57BL6/J mice

Then, we asked whether obesity-elicited CD38^lo^ MRC1^lo^ Macs arising during obesity were derived from monocytes. We used *Ccr2*^−/−^ mice in which the circulating Ly-6C^hi^ monocyte subset mobilizable to sites of inflammation is markedly decreased ^11^. We observed that CD38^lo^ MRC1^lo^ Macs counts were strongly diminished in obese *Ccr2*^−/−^ mice while CD38^hi^ MRC1^hi^ Macs remained similar to controls (Figure 2G). As expected, vWAT Ly-6C^hi^ monocytes counts were lower in obese *Ccr2*^−/−^ mice as compared to controls (Figure 2H). Obesity-elicited CD38^lo^ MRC1^lo^ cells are thus monocyte-derived macrophages.

We showed (Figure 1H and 1I) that recruited CD38^lo^ MRC1^lo^ Macs express higher levels of *Gpnmb, Mmp12, Itgax* and *Il1b* than their CD38^hi^ MRC1^hi^ trMacs counterparts. Accordingly, limited accumulation of CD38^lo^ MRC1^lo^ Macs in obese *Ccr2*^−/−^ mice was associated with decreased mRNA levels of *Gpnmb, Mmp12, Itgax* and *Il1b* in the vWAT as compared to controls (Figure 2I). As previously reported ^2^, obese *Ccr2*^−/−^ mice were more glucose-tolerant than their obese wild-type controls (Figure 2J). Our kinetic analysis revealed that recruited CD38^lo^ MRC1^lo^ Macs accumulation was timely associated with a blunt in vWAT expandability potential (Figure 2F). Along these lines, limited CD38^lo^ MRC1^lo^ Macs infiltration in obese *Ccr2*^−/−^ mice was concomitant with an increase in vWAT mass (Figure 2K), suggesting CD38^lo^ MRC1^lo^ Macs may have a direct impact on vWAT expandability. During obesity, vWAT growth relies on both adipocyte hypertrophy and de novo generation of adipocytes that derive from adipogenic precursors ^12,13^. We recently showed that two main PDGFRα^+^ progenitor subsets differentially expressing CD9 populate the vWAT ^5^. Notably, PDGFRα^+^ CD9^low^ cells demonstrated the unique capacity to differentiate into adipocytes and were gradually lost as obesity develops and vWAT expandability is compromised ^5^. Thus, we asked whether limiting CD38^lo^ MRC1^lo^ Macs infiltration in obese *Ccr2*^−/−^ mice could maintain vWAT expandability by sparing PDGFRα^+^ CD9^low^ pre-adipocytes loss. We indeed observed that PDGFRα^+^ CD9^low^ cells were more numerous in obese *Ccr2*^−/−^ mice compared to controls while PDGFRα^+^ CD9^hi^ progenitors remained unchanged (Figure 2L). This suggests that de novo adipocyte generation favors vWAT growth when CD38^lo^ MRC1^lo^ Macs accumulation is limited.

Altogether, we provide evidence that CD38^lo^ MRC1^lo^ Macs accumulation favors vWAT maladaptive remodeling by fueling inflammation and decreasing PDGFRα^+^ CD9^low^ pre-adipocytes numbers, and both phenomena associated with increased glucose intolerance.

### HFD-induced remodeling of the vWAT macrophage compartment is markedly accelerated in obesity-induced fibrosis prone C3H mice

Most obesity studies are conducted in the C57BL6/J (B6) background, although it does not fully recapitulate the alterations observed in the vWAT of patients with obesity, especially adipose fibrotic remodeling ^4^. Recent studies, including ours, revealed C3H/HeOuJ (C3H) mice are prone to obesity-induced vWAT fibrosis ^5,6^. We thus studied the dynamics of the macrophage subsets in C3H animals compared to B6 mice after 3 and 6 weeks of HFD. As previously described ^5^, vWAT fibrosis was observed as early as 6 weeks of HFD in C3H mice (Figure 3A) and the 3 weeks timepoint thus represents a pre-fibrotic state. In addition, while vWAT growth is gradual in B6 animals over the 6-week period, its expandability potential was readily blunted after 3 weeks of diet in C3H animals (Figure 3B). Regarding macrophage subsets, CD38^hi^ MRC1^hi^ trMacs numbers remained unchanged overtime in both strains (Figure 3C). By contrast, obesity-elicited CD38^lo^ MRC1^lo^ Macs massively accumulated in C3H mice while their numbers marginally increased in B6 animals in these first weeks (Figure 3D). This accumulation of CD38^lo^ MRC1^lo^ Macs was associated with an increased mRNA expression of *Itgax, Mmp12* and *Gpnmb* in the vWAT of HFD-fed C3H mice (Figure 3E), genes we already showed to be preferentially found in CD38^lo^ MRC1^lo^ Macs in B6 animals (Figure 1H). We then sorted CD38^hi^ MRC1^hi^ and CD38^lo^ MRC1^lo^ Macs from lean and obese C3H mice. As observed in B6 animals, CD38^lo^ MRC1^lo^ Macs express higher levels of *Itgax, Gpnmb, Mmp12* and *Spp1* than their CD38^hi^ MRC1^hi^ counterparts (Figure 3F). Thus, as observed at latter time point in B6 animals (Figure 2F), obesity-elicited CD38^lo^ MRC1^lo^ Macs infiltration coincides with the early inability of the vWAT to expand in HFD-fed C3H mice. In addition, CD38^lo^ MRC1^lo^ Macs accumulation also precedes the onset of tissue fibrosis. We then asked whether such remodeling was associated with early changes in progenitor subsets. In agreement with our previous observation, we found that PDGFRα^+^ CD9^low^ pre-adipocytes were already markedly decreased after 3 weeks of HFD, while pro-fibrotic PDGFRα^+^ CD9^hi^ progenitors significantly expanded (Figure 3G). Overall, C3H mice are characterized by extremely rapid and striking switch in the vWAT macrophage populations upon HFD feeding.

**Figure 3.**
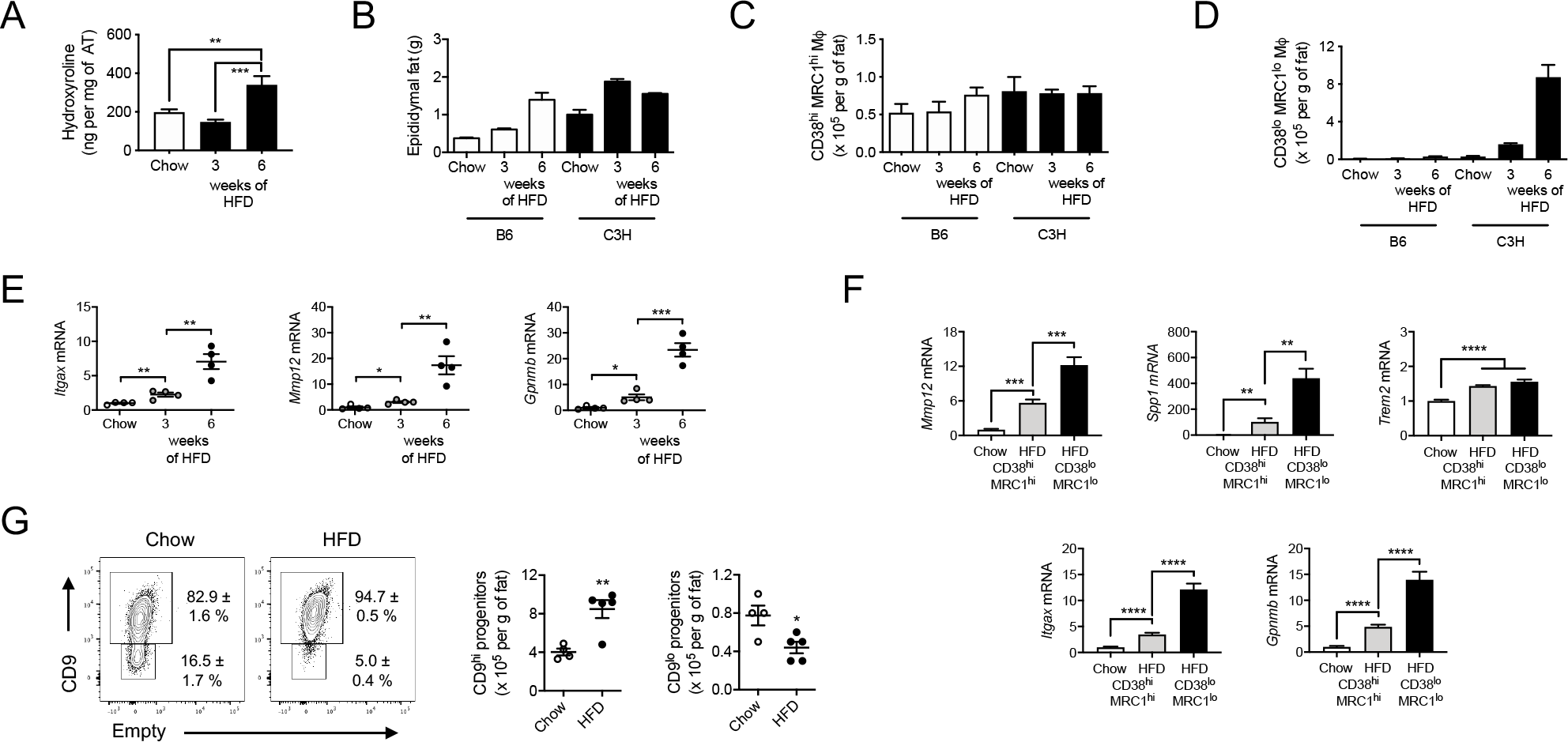
Accelerated obesity-elicited macrophage infiltration associates with adipose tissue fibrosis in obese C3H mice. (A) Hydroxyproline content in the visceral (epididymal) adipose tissue of C3H mice fed a chow diet or a HFD for 3 and 6 weeks (n=5-8 mice per group). (B) Visceral (epididymal) adipose tissue weight in C57BL6/J (B6) and C3H mice before (chow) and after 3 and 6 weeks of HFD (n=3-5 mice per group). (C) CD38^hi^ MRC1^hi^ macrophages in the visceral (epididymal) adipose tissue of C57BL6/J (B6) and C3H mice before (chow) and after 3 and 6 weeks of HFD (n=3-5 mice per group). (D) CD38^lo^ MRC1^lo^ macrophages in the visceral (epididymal) adipose tissue of C57BL6/J (B6) and C3H mice before (chow) and after 3 and 6 weeks of HFD (n=3-5 mice per group). (E) RT-qPCR analysis of monocyte-derived macrophages markers expression in the visceral adipose tissue of C3H mice before (chow) and after 3 and 6 weeks of HFD (n=4 mice per group) (F) RT-qPCR analysis of monocyte-derived macrophage markers expression in CD38^hi^ MRC1^hi^ and CD38^lo^ MRC1^lo^ macrophages sorted from the visceral (epididymal) adipose tissue of C3H mice fed a chow diet or a HFD for 3 weeks (n=6-7 mice per condition). (G) Flow cytometry analysis and quantification of adipose progenitor subsets density in the visceral adipose tissue of C3H mice fed a chow diet or a HFD for 3 weeks (n=4-5 mice per condition).

### Obesity-elicited macrophages fuel vWAT fibrosis in HFD-fed C3H mice

We next wondered whether limiting obesity-elicited Macs accumulation in the vWAT would alter tissue fibrosis. To this aim, we backcrossed C57BL6/J *Ccr2*^−/−^ mice in the C3H/HeOuJ background that sensitizes animals to obesity-induced adipose tissue fibrosis ^5^. As the vWAT of C3H animals undergoes very rapid changes upon HFD feeding, we analyzed C3H *Ccr2*^−/−^ mice and their littermate controls after only 6 weeks of diet, which is sufficient to observed major inflammatory (Figure 3C-D) and fibrotic (Figure 3A) vWAT remodeling. After 6 weeks of HFD, C3H *Ccr2*^−/−^ mice had similar body weight (Figure 4A) and body composition (Figure 4B) than their controls. In addition, vWAT mass (Figure 4C), adipocyte area (Figure 4D) and liver weight (Figure 4E) were also comparable to controls. Flow cytometry analysis of the vWAT revealed a marked decrease in Ly-6C^hi^ monocytes (Figure 4F) and obesity-elicited CD38^lo^ MRC1^lo^ Macs (Figure 4G) in C3H *Ccr2*^−/−^ mice, while trMacs remained similar to controls (Figure 4G). We also assessed progenitors counts by flow cytometry after 6 weeks of diet. At the timepoint, only CD9^hi^ progenitors are found in the vWAT of C3H animals ^5^ and their numbers were significantly lower in the vWAT of C3H *Ccr2*^−/−^ animals as compared to controls (Figure 4H). As CD9^hi^ progenitors are the main extracellular matrix producers and drive the fibrogenic process in the vWAT, we then quantified vWAT fibrosis. We found that vWAT hydroxyproline content was decreased in C3H *Ccr2*^−/−^ mice (Figure 4I). As hydroxyproline accounts for both structural and “fibrotic” collagens, we next stained vWAT sections with Sirius red and imaged the sections under polarized light to specifically quantify collagen networks. Using this method, we observed a marked decrease in fibrosis in the vWAT of C3H *Ccr2*^−/−^ mice as compared to controls (Figure 4J). As extracellular matrix deposition favors vWAT dysfunction and systemic metabolic alterations ^4^, we then assessed the metabolic phenotype of HFD-fed C3H *Ccr2*^−/−^ animals. We first observed that obese C3H *Ccr2*^−/−^ mice were more tolerant to glucose during an oral glucose tolerance test (Figure 4K). In addition, while blood glucose levels were similar in C3H *Ccr2*^−/−^ mice and their controls (Figure 4L), insulin levels were markedly decreased in C3H *Ccr2*^−/−^ animals (Figure 4M). This leads to a lower HOMA-IR index, a systemic indicator of insulin resistance, in C3H *Ccr2*^−/−^ mice (Figure 4N). Finally, we assessed the vWAT response to insulin and showed that insulin-evoked AKT phosphorylation was more pronounced in C3H *Ccr2*^−/−^ mice as compared to controls (Figure 4O). Together, our finding revealed that limiting obesity-elicited CD38^lo^ MRC1^lo^ Macs accumulation in the vWAT of C3H mice decreased CD9^hi^ progenitors, limited fibrosis and improved local and systemic metabolic homeostasis.

**Figure 4.**
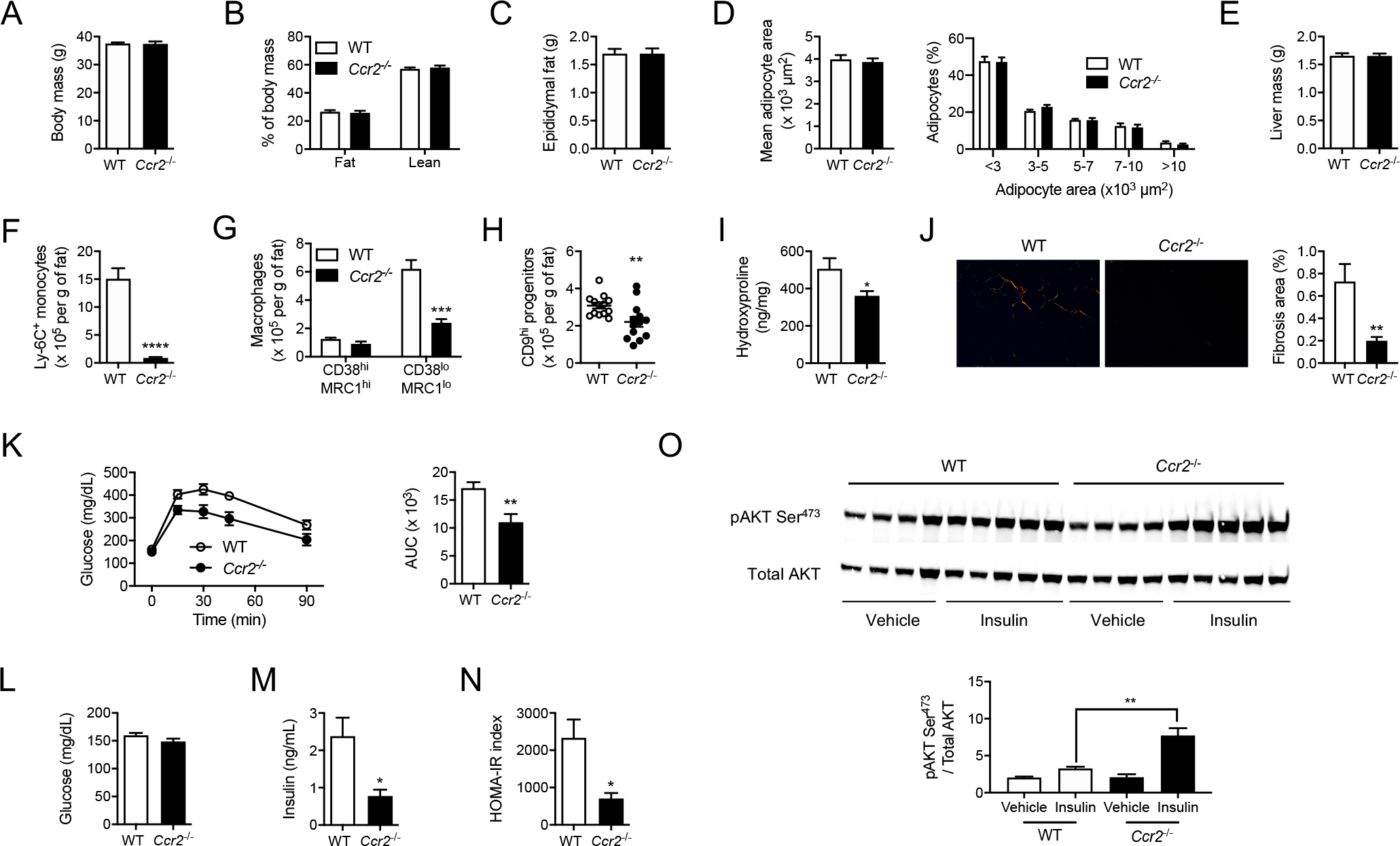
Obesity-elicited macrophage infiltration fuels adipose tissue fibrosis and dysfunction in obese C3H mice. (A) Body weight of C3H *Ccr2*^−/−^ and control animals after 6 weeks of HFD (n=13-17 mice per group). (B) Body composition of C3H *Ccr2*^−/−^ and control animals after 6 weeks of HFD (n=5-7 mice per group). (C) Visceral (epididymal) fat mass in C3H *Ccr2*^−/−^ and control animals after 6 weeks of HFD (n=14-17 mice per group). (D) Mean adipocyte area and adipocyte size distribution in the visceral (epididymal) fat of C3H *Ccr2*^−/−^and control animals after 6 weeks of HFD (n=9-11 mice per group). (E) Liver mass in C3H *Ccr2*^−/−^ and control animals after 6 weeks of HFD (n=14-17 mice per group). (F) Quantification of Ly-6C^+^ monocytes density in the visceral adipose tissue of C3H *Ccr2*^−/−^ and control animals after 6 weeks of HFD (n=7-11 mice per group). (G) Quantification of macrophage subsets density in the visceral adipose tissue of C3H *Ccr2*^−/−^ and control animals after 6 weeks of HFD (n=7-11 mice per group). (H) Quantification of CD9^hi^ progenitors’ density in the visceral adipose tissue of C3H *Ccr2*^−/−^ and control animals after 6 weeks of HFD (n=13 mice per group). (I) Hydroxyproline content in the visceral adipose tissue of C3H *Ccr2*^−/−^ and control animals after 6 weeks of HFD (n=13-16 mice per group). (J) Representative photographs of Sirius red staining under polarized light and quantification of fibrosis area in the visceral adipose tissue of C3H *Ccr2*^−/−^ and control animals after 6 weeks of HFD (n=13-16 mice per group). (K) Glucose tolerance test and quantification of the area under the curve in C3H *Ccr2*^−/−^ and control animals after 6 weeks of HFD (n=8-11 mice per group). (L) Blood glucose levels in C3H *Ccr2*^−/−^ and control animals after 6 weeks of HFD (n=7-10 mice per group). (M) Plasma insulin levels in C3H *Ccr2*^−/−^ and control animals after 6 weeks of HFD (n=7-10 mice per group). (N) HOMA-IR index in C3H *Ccr2*^−/−^ and control animals after 6 weeks of HFD (n=7-10 mice per group). (O) Immunoblot and quantification of AKT (Ser473) phosphorylation status relative to total AKT in the visceral adipose tissue after an intraperitoneal bolus of insulin or vehicle in C3H *Ccr2*^−/−^ and control animals after 6 weeks of HFD (n=4-5 mice per group).

### Obesity-elicited macrophages shape the pro-fibrotic activity of CD9^hi^ progenitors

Our findings reveal that obesity-elicited CD38^lo^ MRC1^lo^ Macs are critical to maintain higher numbers of pro-fibrogenic CD9^hi^ progenitors and to favor vWAT fibrosis. We next sought to determine how these macrophages would impact the function of CD9^hi^ progenitors. First, we performed in-depth analysis of CD38^hi^ MRC1^hi^ trMacs and obesity-elicited CD38^lo^ MRC1^lo^ Macs transcriptional profiles using mRNA sequencing (RNAseq). Macrophage subsets were isolated from the vWAT of chow and HFD-fed C3H animals by cell-sorting. Multiple comparisons were performed to identify differentially expressed genes (DEGs) between cell types and conditions. Then, DEGs were grouped in 6 distinct gene clusters (Figure 5A). The first 3 clusters encompass genes over-represented in trMacs as compared to obesity-elicited Macs. The first cluster comprises genes encoding trMacs markers such as *Pdgfc, Lyve1* or *Folr2*, and includes *Cd38* and *Mrc1* (Figure 5A). Cluster II consists of genes encoding TrMacs markers slightly downregulated in the HFD condition (*Timd4, Cd209a, Cd209b*) (Figure 5A). Cluster III is composed of genes up-regulated in trMacs of HFD-fed animals and contains *Ccl7* and *Ccl12*, which encode ligands of the chemokine receptor CCR2 (Figure 5A). Then, clusters IV and V contain genes over-represented in the HFD conditions. Cluster IV comprises genes markedly induced in trMacs, such as *Csf1*, while cluster V genes were more strongly expressed by obesity-elicited Macs and included *Cd9, Itgax, Mmp12, Osm, Lgals3, Pdgfa* (Figure 5A). Cluster V includes many of the genes that define lipid-associated macrophages (LAMs) ^14^ (*Cd9, Itgax, Mmp12, Gpnmb, Lgals3*). Finally, cluster VI identifies genes over-represented in obesity-elicited Macs and includes, in particular, *Ccr2, Cx3cr1, Il1b, Spp1, Ptgs2, Tnf* and *Pdgfb*. Under HFD, the genes over-represented in trMacs as compared to obesity-elicited Macs were significantly enriched for the Gene Ontology (GO) terms “leukotriene metabolic process” and “lipoxin metabolic process” (Figure 5B). In obesity-elicited Macs, genes associated with the GO terms “cellular response to lipopolysaccharide” and “cytokine-mediated signaling pathway” were strongly enriched (Figure 5C). Overall, moMacs were characterized by their inflammatory signature as compared to trMac. Then, we wondered how limiting obesity-elicited Macs infiltration alters the transcriptional profile of CD9^hi^ progenitors. We thus sorted CD9^hi^ progenitors from HFD-fed C3H *Ccr2*^−/−^ animals and their controls, and analyzed them by RNAseq. The most enriched GO term corresponding to the genes up-regulated in the CD9^hi^ progenitors of *Ccr2*^−/−^ animals was “positive regulation of fat cell differentiation” (Figure 5D), while down-regulated genes were mostly associated with the terms “extracellular matrix organization” and “extracellular structure organization” (Figure 5E). Thus, decreased obesity-elicited Macs numbers dampened the expression of pro-fibrotic genes in CD9^hi^ progenitors, which would explain why adipose tissue fibrosis is decreased in HFD-fed C3H *Ccr2*^−/−^ mice (Figure 4J). To better appreciate the circuitry by which obesity-elicited Macs would impact CD9^hi^ progenitors function, we used the NicheNet ^15^, a computational method that predicts intercellular communications. We exploited this tool to identify the ligands produced by obesity-elicited Macs capable to drive gene expression changes in CD9^hi^ progenitors. The ligands were retrieved from the genes enriched in obesity-elicited Macs as compared to trMacs while receptors were found among the genes expressed by CD9^hi^ progenitors. Finally, we provided a list of DEGs corresponding to genes downregulated in CD9^hi^ progenitors of *Ccr2*^−/−^ animals as compared to wild-type. Indeed, we reasoned that the limited accumulation of obesity-elicited Macs in *Ccr2*^−/−^ animals could contribute to the downregulation of specific genes in CD9^hi^ progenitors. NicheNet revealed several ligand-receptor couples predicted to control gene expression in CD9^hi^ progenitors (Figure 5F and S5). Ligands were mostly linked to GO terms related to cytokine signaling and extracellular matrix organization (Figure 5G). In addition, the target genes predicted to be regulated by the different ligand-receptor couples uncovered were mostly associated with GO terms linked to the organization of the extracellular matrix (Figure 5H). This suggests that obesity-elicited Macs-CD9^hi^ progenitors interactions are linked to fibrogenesis. Interleukin 1β (IL-1β), osteopontin (SPP1), Galectin-3 (GAL3), oncostatin M (OSM), platelet-derived growth factor β (PDGF-BB) and semaphorin-4D (SEMA4D) were among the top ligand candidates, and were predicted to control the expression of key genes associated with fibrogenesis (*Ctgf, Acta2, Fbn1, Serpinb2, Tnc, Col3a1, Timp1, Col1a1, Fn1, Col8a1*, …) (Figure 5F). Thus, obesity-elicited Macs-derived ligands could affect extracellular matrix production and organization through a number of signaling pathways. We then set out to validate whether the NicheNet predictions were valid by treating *in vitro* mouse adipose tissue progenitors with several of the ligands we uncovered above. We assessed both the effect of ligands on cell numbers as well as the acquisition of stress fibers containing α-smooth muscle actin (aSMA) typical of a myofibroblastic phenotype. As a positive control, we validated that the prototypical pro-fibrotic cytokine TGFβ1 heightens both cell counts and αSMA^+^ myofibroblast numbers (Figure 5 I, J). Treatment with IL-1β and GAL3 also increased both parameters (Figure 5 I, J). SEMA4D and PDGFBB only enhanced cell counts while SPP1 only drove αSMA^+^ myofibroblast generation (Figure 5I, J). As progenitors’ numbers and their myofibroblastic activation are key drivers of fibrosis, we combined the two parameters to obtain a fibrotic potential for each ligand (Figure 5L). Together, we revealed that multiple cytokines produced by obesity-elicited Macs can shape progenitors toward a fibrogenic phenotype in the vWAT during obesity.

**Figure 5.**
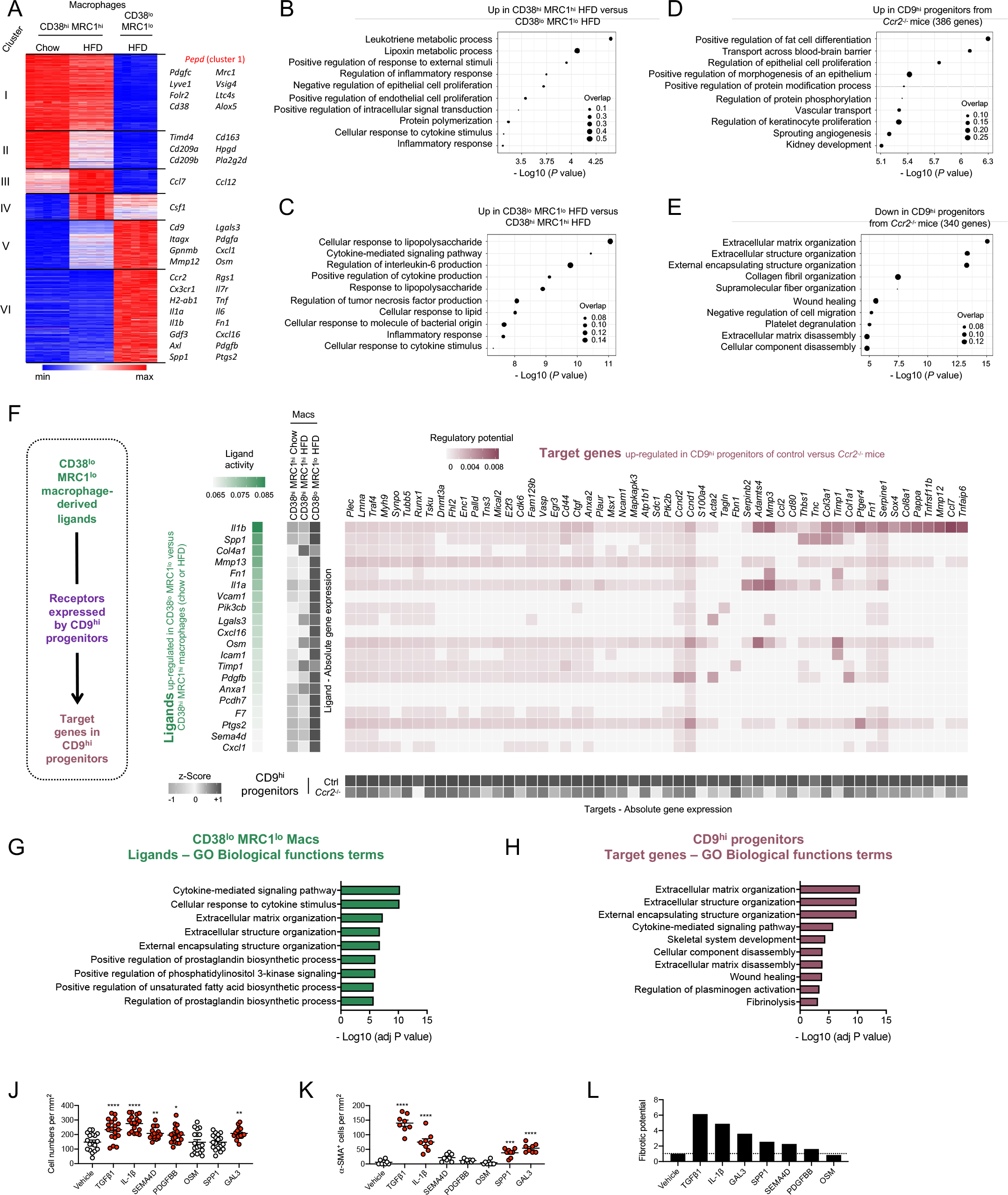
Pathways by which obesity-elicited macrophages impact the phenotype of CD9^hi^ progenitors. (A) K-means clustering analysis of CD38^hi^ MRC1^hi^ and CD38^lo^ MRC1^lo^ macrophages in the visceral WAT of C3H mice fed a chow diet or a HFD for 3 weeks (n=5 mice per group). Representative genes, as well as pathways enriched in those clusters, are also depicted. (B) Pathways enriched in CD38^hi^ MRC1^hi^ macrophages compared to their CD38^lo^ MRC1^lo^ counterparts in HFD fed C3H mice. (C) Pathways enriched in CD38^lo^ MRC1^lo^ macrophages compared to their CD38^hi^ MRC1^hi^ counterparts in HFD fed C3H mice. (D) Pathways enriched in CD9^hi^ progenitors of HFD-fed C3H *Ccr2*^−/−^ mice compared to control animals. (E) Pathways under-represented in CD9^hi^ progenitors of HFD-fed C3H *Ccr2*^−/−^ mice compared to control animals. (F) NicheNet analysis to uncover CD38^lo^ MRC1^lo^ macrophage-derived ligands capable to modulate the phenotype of CD9^hi^ progenitors. (G) Pathways enriched in ligands produced by CD38^lo^ MRC1^lo^ macrophages and predicted to alter gene expression in CD9^hi^ progenitors. (H) Pathways enriched among the genes downregulated in CD9^hi^ progenitors and predicted to be controlled by CD38^lo^ MRC1^lo^ macrophage-derived ligands. (I) Adipose tissue progenitors cell number per mm^2^ after being treated for 24 hours with indicated recombinant proteins. (J) α-smooth muscle actin (α-SMA)-positive adipose tissue progenitors per mm^2^ after being treated for 24 hours with indicated recombinant proteins. (K) Fibrosis potential calculated as the geometric mean of cell number per mm^2^ and α-SMA^+^ progenitors per mm^2^. Geometric means are presented relative to vehicle-treated cells.

### Shift in the macrophage compartment associate with accumulation of pro-fibrotic progenitors and metabolic dysfunctions in patients with obesity

We finally wondered whether the observations made in mice could be translated to human conditions. First, as observed in obese mice, we noticed that the expression levels of *SPP1, ITGAX, TREM2* and *GPNMB* were also elevated in the vWAT of patients with obesity as compared to that of lean controls (Figure 6A). Second, we analyzed human vWAT samples from individuals with obesity by flow cytometry to better characterize the macrophage compartment. We used the pan monocyte/macrophage marker CD14 and found CD14^+^ cells encompass three subsets (Figure 6B). Two subsets, *e*.*g*. CD14^+^ CD206^+^ CD163^+^ CCR2^-^ (subset 1) and CD14^+^ CD206^+^ CD11c^+^ CD163^+^ CCR2^+^ (subset 2) cells, were identified as macrophages (Figure 6C). A third population, CD14^+^ CD11c^+^ CCR2^+^ CD206^-^ CD163^-^ cells, was monocytes (Figure 6C). In addition, the two macrophage subsets separated based on CD209 expression (Figure 6C). Importantly, the differential expression of CD209 and CD11c suggested these subsets may relate to the CD38^hi^ MRC1^hi^ CD209a^hi^ and CD38^lo^ MRC1^lo^ CD11c^+^ Macs observed in mice. We additionally sorted CD209^+^ and CD11c^+^ Macs from human vWAT biopsies and we found that CD11c^+^ Macs express higher mRNA levels of *SPP1, TREM2* and *GPNMB* than their CD209^+^ counterparts (Figure 6D). Altogether, the obese human vWAT contains two Mac subsets with a phenotype resembling the two Mac populations identified in mice. Moreover, we previously reported that, similar to mice, the human vWAT contains two subsets of PDGFRα^+^ progenitors with differential CD9 cell surface expression ^5^. We thus investigated both progenitor and macrophage subsets in the vWAT of patients with obesity and their potential interplay. Interestingly, we here show that the proportion of CD209^+^ Macs was positively associated with the proportion of PDGFRα^+^ CD9^lo^ pre-adipocytes (Figure 6E) while the proportion of CD11c^+^ Macs was positively associated with the proportion PDGFRα^+^ CD9^hi^ fibrogenic progenitors (Figure 6F). Then, in a larger group of patients with obesity, we performed correlation analysis using a set of genes typical of CD11c^+^ Macs (*TREM2, ITGAX, GPNMB* and *SPP1*) that are not expressed in other cell types present in the vWAT (Figure 6G). This transcriptomic signature was positively associated with the proportion of PDGFRα^+^ CD9^hi^ fibrogenic progenitors (Figure 6H, upper panel). We also observed a positive correlation between the frequency of PDGFRα^+^ CD9^hi^ progenitors and IL-1β expression (R=0.399, *P*=0.02; n=31). In addition, the CD11c^+^ Mac transcriptomic signature was positively correlated with the expression of genes encoding key pro-fibrotic mediators (*COL1A1, COL3A1, COL6A1, LOX, CTGF, FN1*) in vWAT biopsies of patients with obesity (Figure 6H, lower panel). This suggests, similar to our observations in mice, that a positive correlation between CD11c^+^ Macs and adipose tissue fibrosis occurs in human obesity. We then analyzed the expression of the CD11c^+^ Mac transcriptomic signature in the vWAT of individuals with obesity stratified as non-diabetic, glucose intolerant and type 2 diabetic. We found a positive association between this signature and blood fasting glucose, fasting insulinemia, HbA1c, the HOMA-IR index of insulin resistance as well as circulating triglycerides (Figure 6I, heat map). Conversely, we found that this transcriptomic signature was negatively correlated with plasma adiponectin levels (Figure 6I, heat map). Circulating C reactive protein (CRP), orosomucoid and interleukin 6 (IL-6) were also evaluated as markers of systemic inflammation, but no association was found with the CD11c^+^ Mac transcriptomic signature (Figure 6I, heat map). Finally, the expression of the CD11c^+^ Mac transcriptomic signature gradually increased with the progressive loss of glucose control in patients with obesity (Figure 6I, bar plot). Overall, our human data suggest that CD11c^+^ Mac accumulation associates with poor glycemic control and shifts the progenitor pool towards its pro-fibrotic fate in agreement with our mouse findings.

**Figure 6.**
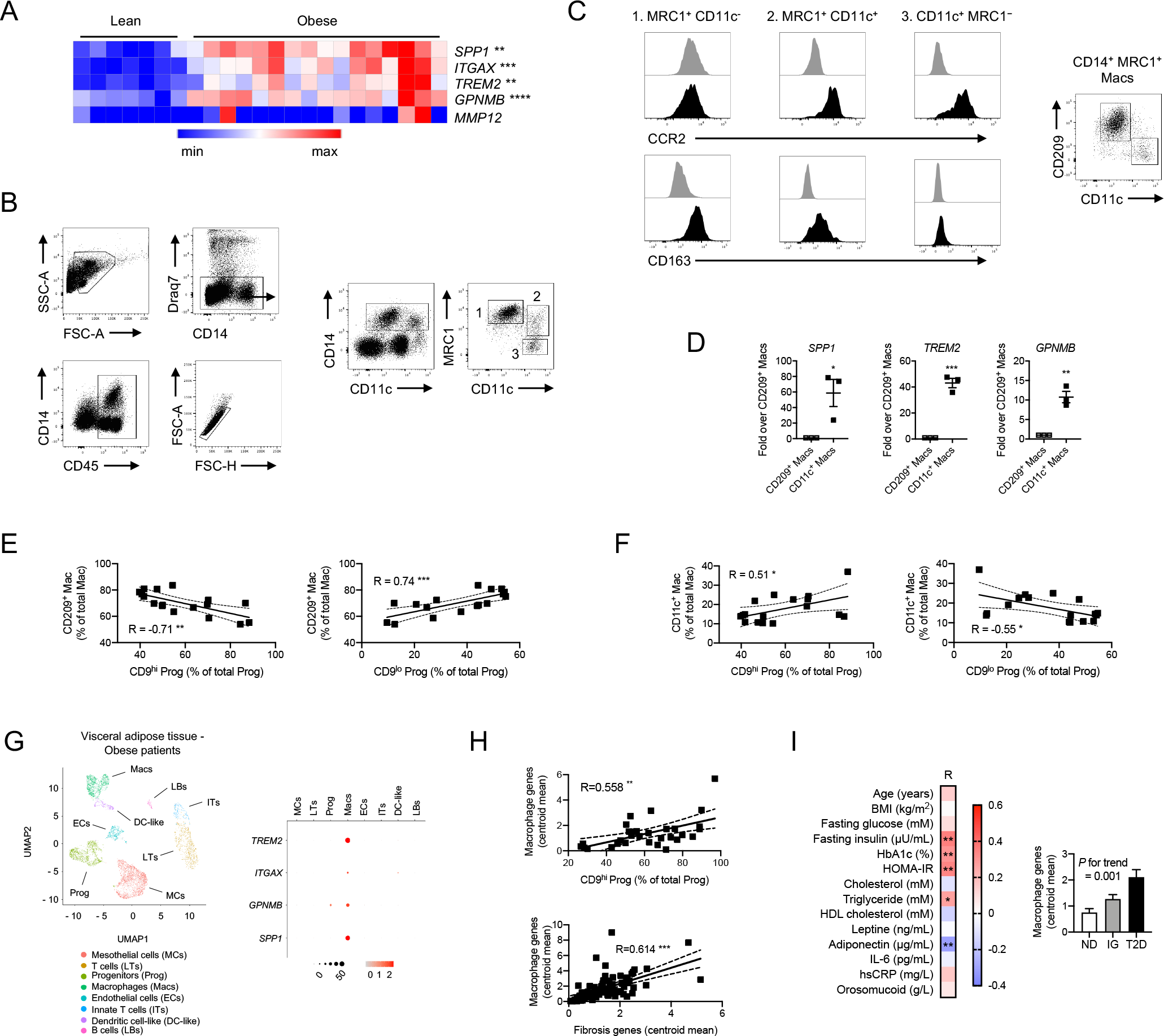
Adipose tissue macrophage and progenitor subsets and their association with clinical parameters in patients with obesity. (A) Microarray analysis of the expression of *SPP1, ITGAX, TREM2, GPNMB* and *MMP12* in the visceral (omental) adipose tissue of lean and patients with obesity obtained from GSE55200. (B) Flow cytometry analysis of the visceral (omental) adipose tissue of patients with obesity. The gating strategy used to define 3 monocyte/macrophage subsets is presented. (C) Expression of the monocyte marker CCR2 and the macrophage marker CD163 by the 3 monocyte/macrophage subsets further helps to discriminate macrophages (MRC1^+^ CD11C^-^ and MRC1^+^ CD11C^+^) from monocytes (CD11C^+^ MRC1^-^). CD209 is expressed by MRC1^+^ CD11C^-^ macrophages but not their MRC1^+^ CD11C^+^ counterparts. (D) RT-qPCR analysis of *SPP1, TREM2* and *GPNMB* expression in CD209^+^ MRC1^+^ CD11C^-^ (CD209^+^ Macs) but not MRC1^+^ CD11C^+^ (CD11C^+^ Macs) (n=3 patients with obesity). (E) Correlation between the frequency of CD209^+^ macrophages and the two adipose progenitor subsets (CD9^hi^ and CD9^lo^) in the visceral (omental) fat of patients with obesity (n=17) (F) Correlation between the frequency of CD11C^+^ macrophages and the two adipose progenitor subsets (CD9^hi^ and CD9^lo^) in the visceral (omental) fat of patients with obesity (n=17) (G) UMAP projection of stromal vascular cells obtained from the visceral (omental) adipose tissue of patients with obesity (obtained from GSE xxx). Dot plot shows the expression of *TREM2, ITGAX, GPNMB* and *SPP1* among the different clusters. (H) Correlation between the centroid mean expression of the CD11C^+^ macrophage markers *TREM2, ITGAX, SPP1* and *GPNMB*, and the frequency of CD9^hi^ progenitors in the visceral (omental) fat of patients with obesity (n=88) (upper panel). Correlation between the centroid mean expression of the CD11C^+^ macrophage markers *TREM2, ITGAX, SPP1* and *GPNMB*, and the centroid mean expression of key pro-fibrotic genes (*COL3A1, COL6A1, LOX, CTGF, FN1* and *INHBA*) in the visceral (omental) fat of patients with obesity (n=88) (lower panel). (I) Heat map showing the correlation between the centroid mean expression of the CD11C^+^ macrophage markers *TREM2, ITGAX, SPP1* and *GPNMB* in the visceral (omental) fat of patients with obesity, and patients’ phenotypic parameters (n=88). The bar plot shows the centroid mean expression of the CD11C^+^ macrophage markers *TREM2, ITGAX, SPP1* and *GPNMB* as a function of patients’ diabetic status (ND: Non-Diabetic, n=15; GI: Glucose Intolerant, n=27; T2D: Type 2 Diabetic, n=45).

## DISCUSSION

Maladaptive vWAT remodeling, which includes both immune and ultrastructural changes, is a key event favoring the onset and progression of obesity comorbidities. Macrophages accumulation and adipose tissue progenitors-mediated fibrosis have been recognized as major events leading to vWAT remodeling and dysfunction ^4,16^. However, it remains unclear whether specific macrophage subsets communicate with adipose tissue progenitors to orchestrate vWAT remodeling and dysfunction during obesity. vWAT macrophage diversity is under intense investigation and we are only beginning to understand how specific subsets alter adipose tissue function. For example, recent work revealed trMacs play a role in adipocyte lipid uptake and hypertrophy ^17^, favoring WAT growth under obesogenic diet ^17,18^. During obesity, WAT inflammation is characterized by Mac accumulation, in particular monocyte-derived Macs. Their role in insulin resistance has been previously reported ^19^ but little is known regarding how these obesity-elicited monocyte-derived Macs control WAT ultrastructural remodeling and whether they shape adipose tissue progenitor fate and function. Furthermore, it remains to determine how results obtained from mice can be translated to human obesity in term of macrophage diversity and vWAT remodeling. In this study, we set out to answer these questions.

We first set out to further characterize macrophages in the obese vWAT of mice. Using a set of markers specific to adipose tissue-resident macrophages, we identified a population of CD38^lo^ MRC1^lo^ CD209a^lo^ monocyte-derived, obesity-elicited macrophages that accumulates over the course of obesity in C57BL6/J mice. Peak accumulation of this macrophage subset concurs with the marked dysregulation of glucose homeostasis. obesity-elicited CD38^lo^ MRC1^lo^ CD209a^lo^ monocyte-derived macrophages express markers previously used to define lipid-associated macrophages in the context of obesity ^14^ as well as scar-associated macrophages in fibrotic liver disease ^20^, suggesting these cells are included in our population. We then assessed macrophage subsets in obese C3H/HeOuj animals with high susceptibility to vWAT fibrosis ^5,6^. We found that obesity-elicited CD38^lo^ MRC1^lo^ Mac accumulation was markedly accelerated in C3H animals as compared to C57BL6/J mice, while CD38^hi^ MRC1^hi^ macrophage numbers were similar in both strains. Thus, remodeling of the macrophage landscape was fast and extreme in the C3H background and future studies will be needed to understand why obesity-elicited Mac accumulation is accelerated in C3H animals as compared to C57BL6/J mice. In addition, this observation questions the use of the C57BL6/J mice as a standard to study HFD-induced obesity. Indeed, as outlined by the hybrid mouse diversity panel, strains responses to a HFD challenge are extremely variable and lead to diverse disease patterns ^21^. Those different patterns may help to better understand the pathogenesis of obesity. We also tried to better define the human adipose tissue macrophage landscape. We found two macrophage subsets defined as CD14^+^ CD206^+^ CD163^+^ CD209^+^ CCR2^-^ (CD209^+^ Macs) and CD14^+^ CD206^+^ CD11c^+^ CD163^+^ CCR2^+^ (CD11c^+^ Macs). Noteworthy, CD11c^+^ Macs share the expression of several markers with mouse CD38^lo^ MRC1^lo^ CD209a^lo^ Macs, suggesting these two subsets might share similar functions. Overall, we found similar macrophage population in the obese vWAT of mouse and humans.

The crosstalk between adipose tissue progenitors and macrophages is starting to be explored. In this context, a recent study revealed that chemokine production by activated adipose tissue progenitors facilitates macrophage accrual in the vWAT of obese animals ^22^. Here, we document how macrophages shape adipose tissue progenitors’ behavior in the vWAT. We have previously shown that CD9^lo^ pre-adipocytes were lost over time in the vWAT of HFD-fed C3H mice ^5^. This was also noticed in C57BL6/J animals although the process took much longer (months versus weeks) ^5^. Here, using HFD-fed *Ccr2*^−/−^ mice in the C57BL6/J background, we observed that CD9^lo^ pre-adipocyte numbers increased when obesity-elicited Mac accumulation was curbed, suggesting these Macs precipitate pre-adipocyte loss during obesity. Alongside, the vWAT mass was increased in HFD-fed *Ccr2*^−/−^ animals and such increased ability to store excess lipids would limit ectopic lipid accumulation and subsequent dysfunction in non-specialized tissues, including the liver. Thus, in the absence of obesity-elicited Macs, adipose tissue progenitors may favor a beneficial expansion of the vWAT through hyperplasia. However, obesity-elicited Macs may also impact on adipocyte triglyceride storage and hypertrophy as mean adipocyte size was previously shown to increase in obese *Ccr2*^−/−^ mice ^19,23^. Thus, future studies will be needed to assess whether adipose progenitors’ participation to the adipocyte pool is increased in *Ccr2*^−/−^ animals. This would need extensive cross-breeding as *Ccr2*-deficient mice will have to be crossed with strains allowing to fate-map progenitors upon HFD feeding, such as *Pdgfrb*-rtTA ^24^ or *Dpp4*-creER^T2 25^ crossed to reporter mice. Overall, obesity-elicited Macs may limit vWAT expandability by controlling both the maintenance of CD9^lo^ pre-adipocytes and adipocyte ability to store lipids. In the C3H background, the loss of vWAT expandability is extremely rapid after the exposition to the HFD as it is already evident after 3 weeks of diet. Similar to C57BL6/J mice, loss of vWAT expansion associates with obesity-elicited Mac accumulation in C3H animals. After 6 weeks of HFD, we did not notice an increase in the vWAT mass in C3H *Ccr2*^−/−^ animals. This could be explained by the late time point studied or by genetic differences. Thus, limiting obesity-elicited Mac buildup helped maintain the pre-adipocyte pool and this was associated with increased vWAT expandability in C57BL6/J mice. Next, we sought to study how obesity-elicited Macs alter CD9^hi^ progenitors’ behavior, especially their role in ECM production and the development of vWAT fibrosis. To this aim, we backcross *Ccr2*^−/−^ animals in the fibrosis-prone C3H background ^5,6^. Indeed, In C3H mice, analysis of two time points characterizing pre-fibrotic and fibrotic stages revealed that obesity-elicited Mac accumulation was readily observable in the pre-fibrotic vWAT before tissue collagen deposition occurs, suggesting they may be integral to the fibrotic process. Obesity-elicited Mac numbers were dampened in the vWAT of C3H *Ccr2*^−/−^ mice and this was associated with decreased tissue fibrosis. CD9^hi^ progenitors are the main drivers of HFD-induced vWAT fibrosis and their numbers increase in response to HFD feeding in C3H animals ^5^. We observed that this increase was limited in the absence of obesity-elicited Macs. In addition, we showed that genes linked to GO terms related to fibrogenesis were downregulated in CD9^hi^ progenitors sorted from HFD-fed C3H *Ccr2*^−/−^ mice as compared to control animals. Thus, we showed that limiting obesity-elicited Mac buildup in C3H mice impaired PDGFRα^+^ CD9^hi^ progenitors’ expansion and their engagement towards fibrogenesis, leading to decreased vWAT fibrosis and improve glucose tolerance.

In humans, we found that CD14^+^ CD206^+^ CD11c^+^ CD163^+^ CCR2^+^ (CD11c^+^ Macs) have a similar signature than mouse obesity-elicited macrophages. CD11c^+^ Macs were positively associated with the proportion of CD9^hi^ progenitors in the vWAT of obese patients. In addition, a CD11c^+^ Mac signature was positively correlated with increased CD9^hi^ progenitors, vWAT fibrosis and impaired glucose control in obese patients. Thus, we identified in patients with obesity a macrophage population related to mouse obesity-elicited Mac that is similarly linked to vWAT fibrosis. These findings suggest that targeting CD11c^+^ Macs hold promise to curb vWAT fibrosis and dysfunction in humans.

We uncovered a crosstalk between obesity-elicited macrophages and CD9^hi^ adipose tissue progenitors. We thus sought to determine how obesity-elicited macrophages shape the pro-fibrogenic activation of CD9^hi^ progenitors. We used RNAseq of specific cell populations and a cell-cell communication analysis tool (NicheNet) to answer this question. The NicheNet algorithm identified several ligands predicted to signal on CD9^hi^ progenitors and control pro-fibrotic gene expression. Among predicted ligands, IL-1β, SPP1, GAL3, SEMA4D and PDGF-BB were shown to control progenitors’ proliferation and/or differentiation toward myofibroblasts. Thus, a number of ligands secreted by obesity-elicited macrophages can command the fibrogenic activation of CD9^hi^ progenitors. This suggests that vWAT fibrosis is controlled by a complex network of macrophage-secreted mediators that would act in concert to increase progenitors numbers and ECM production capacities, leading to elevated ECM deposition and organization.

In summary, our study further describes the diversity of adipose tissue macrophages in obesity and revealed that monocyte-derived, obesity-elicited macrophages impact vWAT remodeling by modulating the fate and function of adipose tissue progenitors in mice and humans. We more particularly revealed how obesity-elicited macrophages shape the pro-fibrogenic activation of CD9^hi^ progenitors. Our findings thus provide a framework to develop strategies aimed at limiting vWAT maladaptive remodeling and associated local and systemic metabolic alterations in obesity.

## METHODS

### Mouse strains used in the study

C57BL/6J mice were from Janvier Labs or bred in house. C3H mice were obtained from Charles River or bred in house. *Ob/Ob* (B6.Cg-*Lep*^ob^/J) mice were from Charles River. *Ccr2*^−/−^ (B6.129S4-*Ccr2*^*tm1Ifc*^/J) mice were obtained from the Jackson Laboratory and backcrossed in-house to C3H mice. We found C3H propensity to develop adipose tissue fibrosis in response to HFD feeding is a dominant trait in F1 (50% of C57BL/6J alleles; 50% of C3H alleles) animals. Here, *Ccr2*^−/−^ animals with a minimum of 5 backcross generations to the C3H background strain (N5 *i*.*e*. 96.9% of C3H alleles) have been studied.

### Animal housing and diets used

Mice were maintained on a 12-hour light and dark cycle with ad libitum access to water and standard chow diet (no. 5058; Lab-Diet). For obesity studies, animals were fed a high-fat diet (HFD, 60% calories from fat, D12492, Research Diets). Diet duration was indicated in the text and/or figure legends. All animal procedures were reviewed and approved by local and national comities, and conducted in accordance with the Guide for the Care and Use of Laboratory Animals published by the European Commission Directive 86/609/EEC.

### Human subjects

Omental WAT samples were obtained from subjects with obesity enrolled in the bariatric surgery program at the Nutrition Department of Pitié-Salpétrière hospital in Paris. Ethical approval was obtained from the Research Ethics Committee of Hôtel-Dieu Hospital (CPP Ile-de-France N°1). Informed written consent was obtained from all subjects and the protocol was registered on http://www.clinicaltrials.gov (NCT01655017, NCT00476658). The clinical characteristics of the subjects used for Figure 4I are described in Table S1. Before BS, height and weight were measured by standard procedures. Blood samples were collected after a 12 hours overnight fast and clinical variables were measured as described ^28^. Glucose intolerance and type 2 diabetes were diagnosed in accordance with the American Diabetes Association definition.

### Fat and lean mass measurement

Fat and lean mass were measured by TD-NMR using a MinispecPlus LFII90 body composition analyzer (Bruker).

### Glucose metabolism assessment

For assessment of oral glucose tolerance, mice were fasted for 5 hours prior to intraperitoneal glucose injection at a dose of 2 grams per kg of body weight. Glycaemia was measured with a glucometer (Accu-Check, Roche) at baseline and 15, 30, 60, 90 and 120 minutes after gavage. Blood was also collected at baseline for insulin determination. Insulin levels were measured with the mouse ultrasensitive insulin ELISA kit from Alpco. The HOMA-IR index, an indicator of insulin resistance, was calculated with the following formula: fasting plasma insulin (mU/mL) × fasting plasma glucose (mm/L)/22.5. To assess insulin resistance *in situ*, animals were injected with insulin (8 units per kg) or vehicle intraperitoneally and their visceral adipose tissue (epididymal depot) was collected 8 minutes later. Tissues were processed to isolate proteins and immunoblots of AKT (total and phosphorylated at Ser473) were performed as previously described ^5,29^

### processing and cell suspension preparation

Mouse epidydimal adipose tissue was harvested and digested in Hanks’ Balanced Salt Solution (HBSS, no calcium and magnesium) containing fetal bovine serum (FBS,3%) and collagenase D (2.5 mg/ml, Sigma-Aldrich) for 30 minutes at 37°C under agitation. After homogenization, the suspension is washed in PBS buffer containing bovine serum albumin (1%) and EDTA (0.5%). After centrifugation, the adipocyte fraction is discarded and the stroma vascular fraction passed through a 100 µm filter before staining.

Human omental adipose tissue was digested in DMEM medium containing bovine serum albumin (1%), HEPES (0.1M) and collagenase A (1 mg/l), and then processed similarly to mouse samples.

### Flow cytometry

Mouse and human antibodies were purchased from Biolegend, Thermo Fisher Scientific, Miltenyi Biotec and BD Biosciences. For mice, the following markers and clones were used: CD115 (AFS98), CD11c (N418), MHC-II (M5/114.15.2), CD11b (M1/70), CD45 (30-F11), Gr-1 (RB6-8C5), Ly-6C (HK1.4), CD64 (X54-5/7.1), F4/80 (BM8), Ki-67 (B56), SIGN-R1 (CD209b, REA125), LYVE1 (ALY7), Siglec-F (E50-2440), CD206 (C068C2), CD301 (LOM-14), CD209a (MMD3), CD38 (90), CD31 (390), CD140a (APA5), PDPN (8.1.1) and CD9 (MZ3). For human studies, the following markers and clones were used: CD140A (aR1), CD9 (eBioSN4), CD34 (581), CD45 (5B1 ou H130), CD31 (WM59), CD14 (61D3), CCR2 (K036C2), CD163 (GHI/61.1), CD206 (DCN228), CD11C (Bu15) and CD209 (DCN47.5).

Cell suspensions were stained with appropriate antibodies for 30 min on ice. Draq7 or propidium iodide were used to exclude dead cells. Intracellular Ki-67 staining was performed using the Foxp3 staining kit from Thermo Fisher Scientific. Bodipy staining was performed on cells fixed using the Cytofix/Cytoperm™ kit from BD Biosciences. Lysotracker Red staining was performed accordingly to the manufacturer specifications.

When absolute cell counts are provided, a fixed number of nonfluorescent beads (10000, 10-µm polybead carboxylate microspheres from Polysciences) was added to each tube. The formula number of cells = (number of acquired cells × 10,000) / (number of acquired beads) was used. Cell counts were finally expressed as a number of cells per milligram of tissue. Data were acquired on a BD LSRFortessa™ flow cytometer (BD Biosciences) and analyzed with FlowJo software (Tree Star). Cell sorting was performed on a BD FACSAria II™ cell sorter.

### Biochemical measurement in mice

Plasma alanine aminotransferase (ALT) levels were determined using a UV test kit (ALAT GPT FS, Diasys) run on a Konelab analyzer (Thermo Fisher Scientific).

### mRNA extraction, reverse transcription and quantitative PCR

RNAs from epidydimal adipose tissue samples were prepared using the RNeasy Lipid Tissue kit (Qiagen) and cDNAs were synthesized using the Transcriptor First Strand cDNA Synthesis Kit (Roche).

RNAs from sorted cells were prepared using the RNeasy Plus micro kit (Qiagen). RNA was reverse transcribed using the SuperScript VILO cDNA synthesis kit (Thermo Fisher Scientific). Quantitative PCR analyses were performed using a LightCycler® 480 real-time PCR system and dedicated software (Roche). Gene expression was normalized to a minimum of 2 housekeeping genes using the Roche LightCycler® 480 software. All primer sequences are available upon request.

### Hydroxyproline content

Hydroxyproline content was measured using a hydroxyproline colorimetric assay (BioVision). Briefly, frozen epidydimal fat is weighted, distributed in sealed tubes containing 6N HCl (10 μL of HCl/mg of WAT) and heated overnight at 110°C. Ten microliters are evaporated before incubation with chloramine-T and p-di-methyl-amino-benzaldehyde (DMAB) at 60°C for 90 min. The absorbance is read at 560 nm and the concentration is determined using the standard curve generated with purified hydroxyproline.

### Histological analysis of the liver and adipose tissue

Epidydimal tissue and liver samples are fixed in 10% formalin buffer and then embedded in paraffin to produce tissue sections (5 µm). Liver sections were stained with hematoxylin and eosin to identify steatotic areas. Epidydimal adipose tissue collagen content was determined by staining sections with picrosirius red (Sigma-Aldrich) and photographs are taken under polarized light to reveal crosslinked collagens. For quantitation of adipocyte size, adipose tissue sections were stained with hematoxylin and eosin, and the Calopix software was used to measure average adipocyte area as well as the distribution of adipocyte sizes.

### Publicly available microarray and single-cell mRNA sequencing (scRNAseq) datasets

Epidydimal adipose tissue microarray data of lean and obese male mice (GSE36033, in which mice were fed with the same high-fat diet than in our current study) ^30^ were used to obtain a signature of the genes induced in the vWAT during obesity. We also used our previous microarray profiling of sorted mouse blood monocytes and thioglycolate-elicited peritoneal monocyte-derived macrophages (GSE15907) ^10^ to obtain a gene signature typical of monocyte to macrophage differentiation *in vivo*. Finally, microarray profiling of omental adipose tissue samples from patients with obesity and lean controls (GSE55200) ^31^ were used to determine the expression levels of several genes of interest.

Human omental adipose tissue scRNAseq data were obtained from a published study ^32^ and analyzed using the Seurat R package.

### mRNA sequencing, data preprocessing and analysis

Total RNA preparation was performed from 20000 sorted cells (macrophage subsets or adipose tissue CD9^hi^ progenitors) using the RNeasy Plus Micro Kit (Qiagen). cDNA libraries were generated using the KAPA mRNA HyperPrep kit (Roche). RNA-Seq libraries were sequenced on an Illumina NovaSeq 6000 (30 million reads per sample). RNAseq analysis was completed using the Eoulsan 2.5 pipeline. The minimap2 version 2.18 index was used to map raw reads to the genome and data normalization was performed with DESeq2. A linear model for microarray and RNAseq analysis (LIMMA) was conducted to select differentially expressed genes with a 1.5-fold change cutoff between at least two conditions. Annotated genes with a count mean over 10 in at least one condition and a coefficient of variation of more than 0.5 between at least two conditions were retained. Principal component analysis, K-means clustering and GSEA analysis were performed using R. K-means clustering and cluster number determination were performed using R and visualized using the Phantasus web platform. The R package Enrichr was used for pathway analysis and illustration. Data were deposited at NCBI under the accession number E-MTAB-12596.

### NicheNet analysis

To analyze cell-cell communications, we applied the NicheNet algorithm that integrates knowledge on ligand-to-target signaling paths ^15^. Using gene expression data of macrophage subsets and CD9^hi^ progenitors as inputs, we aimed to uncover how macrophages could impact on CD9^hi^ progenitors’ behavior by predicting the ligand-receptor interactions that modulate gene expression in CD9^hi^ progenitors. “Sender” genes (ligands) were identified among the genes induced (minimum of 2-fold change) in CD38^lo^ MRC1^lo^ Macs as compared to CD38^hi^ MRC1^hi^ Macs in the chow and HFD conditions. “Receivers” (receptors) were identified among the genes expressed by CD9^hi^ progenitors. Once the ligand-receptors (L-R) pairs are identified, the NicheNet algorithm looks for potential gene targets of these L-R pairs in a defined “geneset”. L-R pairs targets are integral to the algorithm and consist of a prebuiltprior model that integrates existing knowledge on ligand-to-target signaling pathways. Here, the “geneset” comprised genes that were downregulated (with a minimum of 25% decrease in gene expression) in CD9^hi^ progenitors of *Ccr2*^−/−^ mice that present a marked decrease in CD38^lo^ MRC1^lo^ Macs accumulation in their vWAT. Once run, Nichenet returns a list of prioritized ligands with the target genes they are predicted to control. All analyses were conducted according to the NicheNet tutorial (https://github.com/saeyslab/nichenetr).

### Analysis of mouse adipose progenitors proliferation and myofibroblastic differentiation *in vitro*

Epidydimal adipose tissue from lean mice was processed as described above to obtain the non-adipocyte fraction. Cells were seeded in 75cm^2^ flask and cultured in DMEM supplemented with 10% FBS. After two passages, progenitors were seeded in Lab-Tek 16-well glass slides (Thermo Fisher Scientific). After reaching confluence, cells were placed in serum free-DMEM for 24 h prior to stimulation with either TGFβ1 (1ng/ml) or IL-1β, PDGFBB, SPP1, OSM, GAL3 or SEMA4D (all at 10ng/ml and obtained from R and D Systems) for 24h. Cells were then fixed in formalin, permeabilized in PBS-0.1% TritonX-100 for 4 min at room temperature (RT) and blocked in PBS-0.25% fish gelatin for 30 min at RT. Next, cells were successively incubated with the primary mouse anti-human αSMA antibody (clone 1A4) and then with the secondary Alexa Fluor 594-conjugated anti-mouse antibody. Slides were mounted with fluoromount containing DAPI (Vector Laboratories). Images were captured using a Zeiss Axio Imager M2 fluorescence microscope and analyzed using FIJI software.

### Statistical analysis

Statistical significance of differences was performed using GraphPad Prism (GraphPad Software). Two-tailed Student’s t-test was used to assess the statistical significance of the difference between means of two groups. When more than 2 groups were studied, analysis of variance was performed followed by Newman-Keuls as post-hoc test. Graphs depicted the mean ± SEM. Statistical significance is represented as follows: *P<0.05, **P<0.01, ***P<0.001 and ****P<0.0001.

## AUTHOR CONTRIBUTIONS

Conceptualization, E.L.G. and G.M.; methodology, E.L.G., G.M., K.C., and T.H.; investigation, H.A., S.D., M.C.d.O., A.O., L.P., C.R., M.O., R.M., M.M., G.M. and E.L.G.; writing – original draft, E.L.G. and G.M.; writing – review and editing, A.C., T.H., K.C., G.M. and E.L.G; supervision, E.L.G. and G.M.; funding acquisition, E.L.G., K.C. and G.M.

## ACKNOWLEDGMENTS

This work was supported by grants to ELG from the Agence Nationale pour la Recherche (ANR-17-CE14-0023, ANR-17-CE14-0009, ANR-21-CE14-0023). GM and KC received funding from the Agence Nationale pour la Recherche (ANR-17-CE14-0009). GM received funding from the Agence Nationale pour la Recherche (ANR-21-CE14-0023) and the European Foundation for the Study of Diabetes (EFSD). Melissa Ouhachi received a one-year doctoral fellowship from the Nouvelle Société Française d’Athérosclérose (NSFA). Marina Chaves de Oliveira and Amanda Oliveira were supported by the Coordenação de Aperfeiçoamento de Pessoal de Nível Superior (CAPES, Brazil).

