## Supplemental figures for "Obesity-elicited macrophages shape CD9^hi^ progenitor fate to promote adipose tissue fibrosis and dysfunction"

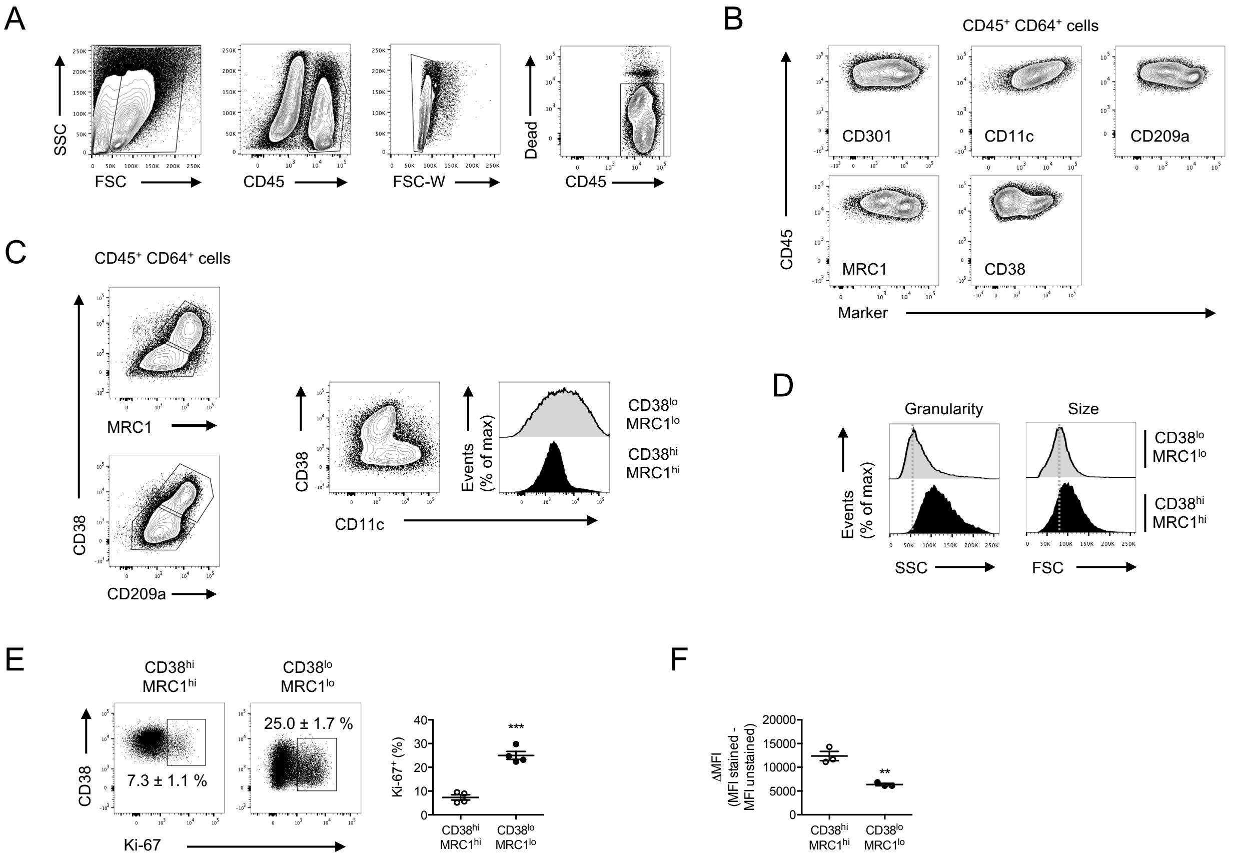


**Figure S1. Identification of two major macrophage subsets in the adipose tissue of obese mice.**

(A) Gating strategy to study live $CD45^+$ leucocytes in the stromal vascular fraction of the vWAT.

(B) Cell surface expression of CD301, CD11c, CD209a, MRC1 and CD38 by $CD64^+$ vWAT macrophages of obese animals.

(C) The expression of CD38, MRC1 and CD209a defines two macrophage subsets in the vWAT of genetically-obese *Ob*/*Ob* animals.

(D) Granularity and size of $CD38^{hi}$ $MRC1^{hi}$ and $CD38^{lo}$ $MRC1^{lo}$ macrophages in the vWAT of obese mice.

(E) Expression of the proliferation marker Ki-67 by $CD38^{hi}$ $MRC1^{hi}$ and $CD38^{lo}$ $MRC1^{lo}$ macrophages in the vWAT of obese mice (n=4 mice).

(F) Quantification of intracellular lipid levels by Bodipy in $CD38^{hi}$ $MRC1^{hi}$ and $CD38^{lo}$ $MRC1^{lo}$ vWAT macrophages of obese mice (n=3 mice).

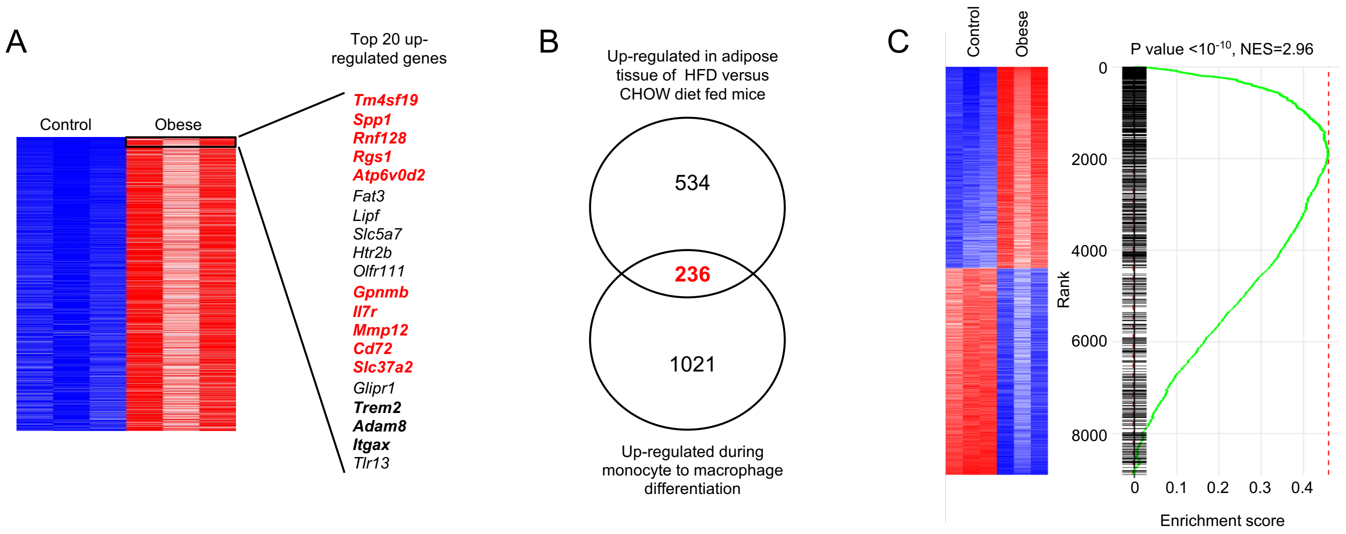


**Figure S2. The obese adipose tissue gene signature is dominated by transcripts associated with monocyte to macrophage differentiation.**

(A) Representation of the top 20 genes up-regulated in the vWAT of HFD-induced obese animals as compared to chow-fed lean controls (obtained from GSE36033)

(B) Venn diagram showing the overlap (236 genes, including those highlighted in red in panel A) between genes up-regulated in the vWAT of obese mice (n=770) and genes induced during monocyte to macrophage differentiation (n=1257)

The monocyte-derived macrophage gene signature was obtained from our previous study on thioglycolate-elicited macrophages (GSE15907)

(C) Gene Set Enrichment Analysis (GSEA) of the monocyte to macrophage differentiation gene signature (1257 genes) in genes up- and down-regulated in the vWAT of HFD-induced obese animals as compared to chow-fed lean controls.

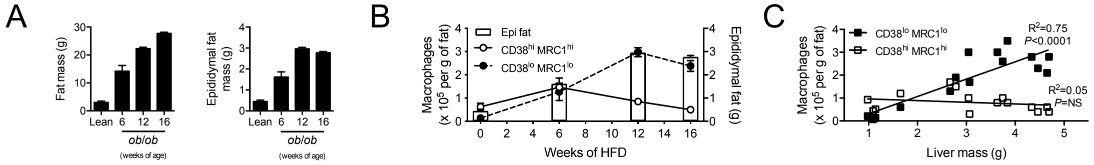


**Figure S3. Loss of visceral fat expandability associates with obesity-associated CD38$^{lo}$ MRC1$^{lo}$ macrophages accumulation in genetically-obese *Ob*/*Ob* mice.**

(A) Total body fat mass determined by nuclear magnetic resonance and visceral (epidydimal) fat weight in 6, 12 and 16 weeks-old genetically obese *Ob*/*Ob* mice and 6 weeks-old lean controls (n=3-5 per group).

(B) Macrophage subsets density (lines) and visceral (epididymal) fat mass (bar) over the time course of HFD-induced obesity in 6, 12 and 16 weeks-old genetically obese *Ob*/*Ob* mice and 6 weeks-old lean controls (n=3-5 per group).

(C) Correlation between macrophage subsets density and liver mass using *Ob*/*Ob* animals at different ages and lean controls (n=15 mice total).

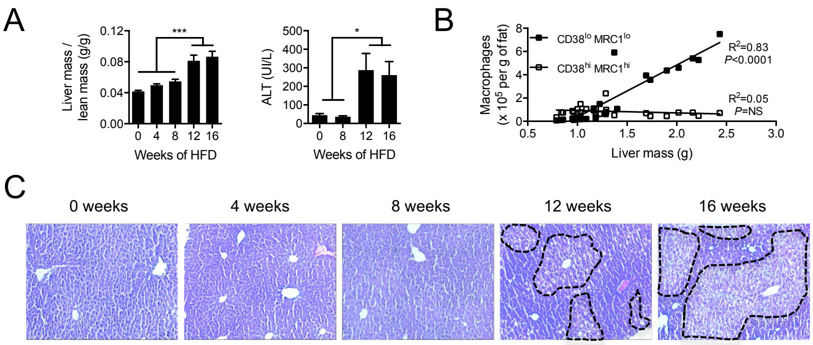


**Figure S4. Loss of visceral fat expandability associates with liver alteration in HFD-fed animals.**

(A) Liver to lean mass ratio over the course of HFD-induced obesity (n=5 per time point).

(B) Plasma transaminase levels over the course of HFD-induced obesity (n=4 per time point).

(C) Photographs of representative liver sections over the course of HFD-induced obesity. Dashed lines surround areas of steatosis.

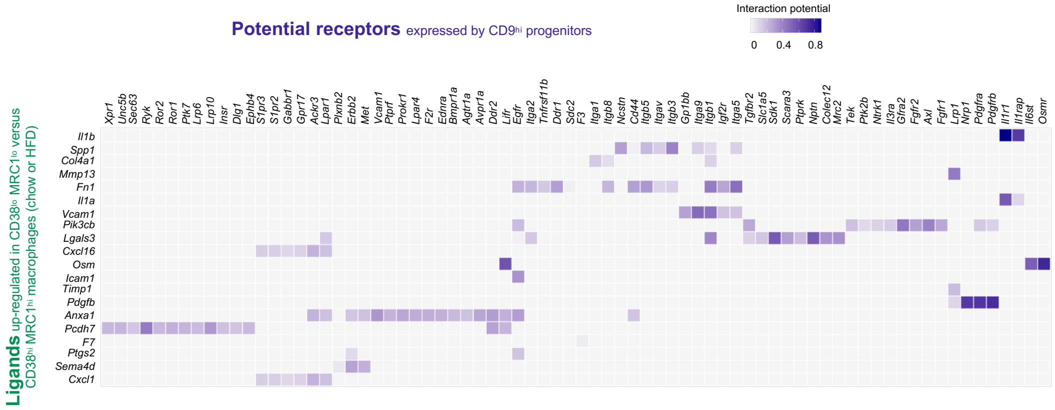


**Figure S5. Receptors expressed by CD9$^{hi}$ progenitors that can bind to the ligands secreted by obesity-associated CD38$^{lo}$ MRC1$^{lo}$ macrophages.**

This heatmap represent the ligand-receptors pairs identified by the NicheNet algorithm

| Clinical parameters | Obese (n=18) | Obese GI (n=27) | Obese T2D (n=49) |
|---|---|---|---|
| Female, n (%) | 15 (83) [ab] | 25 (93) [a] | 31 (63) [b] |
| Age, year | 33.28 ± 2.3 [a] | 46.74 ± 2.3 [b] | 49.47 ± 1.8 [b] |
| BMI, kg/m$^2$ | 43.87 ± 1.6 | 48.11 ± 1.5 | 43.96 ± 1.1 |
| Fasting glycemia, mM | 4.82 ± 0.1 [a] | 5.32 ± 0.1 [a] | 8.06 ± 0.3 [b] |
| Fasting insulin, mU/ml | 15.82 ± 2.3 | 20.06 ± 2.6 | 19.74 ± 1.5 |
| HbA1c, % | 5.32 ± 0.0 [a] | 5.88 ± 0.0 [a] | 7.73 ± 0.2 [b] |
| HOMA-IR | 3.37 ± 0.4 [a] | 4.72 ± 0.6 [a] | 6.87 ± 0.6 [b] |
| Total cholesterol, mM | 4.95 ± 0.2 | 4.92 ± 0.1 | 4.56 ± 0.1 |
| Triglycerides, mM | 1.24 ± 0.1 [a] | 1.28 ± 0.1 [a] | 1.91 ± 0.1 [b] |
| HDL cholesterol, mM | 1.30 ± 0.1 [a] | 1.27 ± 0.1 [a] | 1.05 ± 0.0 [b] |
| Adiponectin, mg/ml | 5.51 ± 0.6 [a] | 4.55 ± 0.4 [a] | 3.29 ± 0.2 [b] |
| Leptin, ng/ml | 60.66 ± 7.3 [ab] | 72.08 ± 6.1 [a] | 49.32 ± 3.4 [b] |
| hsCRP, mg/L | 6.74 ± 1.1 | 10.29 ± 1.5 | 7.65 ± 0.7 |
| IL-6, pg/mL | 3.73 ± 0.5 | 5.24 ± 0.5 | 4.75 ± 0.4 |
| Orosomucoid, g/L | 0.91 ± 0.1 | 0.97 ± 0.0 | 0.88 ± 0.0 |

**Table S1. Clinical parameters of the patients studied in Figure 4I.**
Kruskal-Wallis analysis and Dunn's multiple comparison test were performed. For categorical data, Fisher's exact test was used. Different letters (a, b) show statistical significance (*P*<0.05).
